# Thetis cells induce food-specific Treg cell differentiation and oral tolerance

**DOI:** 10.1101/2024.05.08.592952

**Authors:** Yollanda Franco Parisotto, Vanja Cabric, Tyler Park, Blossom Akagbosu, Zihan Zhao, Yun Lo, Logan Fisher, Gayathri Shibu, Yoselin A. Paucar Iza, Christina Leslie, Chrysothemis C. Brown

## Abstract

The intestinal immune system must establish tolerance to food antigens to prevent onset of allergic and inflammatory diseases. Peripherally generated regulatory T (pTreg) cells play an essential role in suppressing inflammatory responses to allergens; however, the antigen-presenting cell (APC) that instructs food-specific pTreg cells is not known. Here, we show that antigen presentation and TGF-β activation by a subset of RORγt^+^ antigen-presenting cells (APC), Thetis cells IV (TC IV), is required for food-induced pTreg cell differentiation and oral tolerance. By contrast, antigen presentation by dendritic cells (DCs) was dispensable for pTreg induction but required for T_H_1 effector responses, highlighting a division of labor between tolerogenic TCs and pro-inflammatory DCs. While antigen presentation by TCs was required for food-specific pTreg generation both in early life and adulthood, the increased abundance of TCs in the peri-weaning period was associated with a window of opportunity for enhanced pTreg differentiation. These findings establish a critical role for TCs in oral tolerance and suggest that these cells may represent a key therapeutic target for the treatment of food-associated allergic and inflammatory diseases.

---

The immune system must establish tolerance to orally ingested food proteins to ensure intestinal homeostasis. Failure to do so results in onset of allergic or inflammatory diseases. Peripherally induced Foxp3^+^ regulatory T (pTreg) cells are instrumental in suppressing inflammatory responses to dietary antigens^1–4^; however, the mechanisms underlying the development of food-specific Treg cells are not well understood. The current paradigm is that dendritic cells (DC) induce pTreg cell differentiation and oral tolerance^5^; however, to date, the ‘tolerogenic’ DC subset responsible for pTreg induction has not been identified.

Although oral tolerance has been widely studied in adult mice, dysregulated responses to food antigens including IgE-mediated food allergy, food protein-induced inflammatory enterocolitis and celiac disease, present almost exclusively in infancy or childhood, when food antigens are first encountered. Commensurate with a developmental window for oral tolerance, early introduction of allergenic foods, prior to the WHO-recommended age of 6 months, is associated with reduced incidence of food allergy^6–9^. Moreover, oral immunotherapy trials in food allergic children have highlighted a window of opportunity for therapeutic benefit^10^. However, the cellular and molecular mechanisms underpinning the increased propensity for oral tolerance in early life are not known.

We hypothesized that layered ontogeny of tolerogenic antigen-presenting cells ensures tolerance to dietary antigens during the period in which they are first encountered. We recently identified a novel lineage of RORγt^+^ APCs, Thetis cells (TC), enriched in gut lymph nodes during early life. These cells included a subset of cells (TC IV) that induce pTreg differentiation and tolerance to the gut microbiota^11–14^ suggesting a potential role for TC IV in tolerance to oral antigens.

## Thetis cells induce food-specific pTreg cells and oral tolerance

To establish the full spectrum of antigen-presenting cells (APC) that present food antigens during the weaning period, we administered ovalbumin (OVA) conjugated to AF647 intragastrically (i.g.) to postnatal day 14 (P14) mice and analyzed antigen uptake by MHCII^+^ cells in small intestinal mesenteric lymph nodes (siLN) 16 h later. This revealed predominant uptake by two APC types: CCR7^+^ (migratory) type 1 dendritic cells (DC1s) and RORγt^+^MHCII^+^CXCR6^−^CD11b^+^ TC IV, as well as a minor fraction of CCR7^+^ DC2s (**Fig. 1, A and B**, and **fig. S1A**). RORγt^+^ APCs expressing Itgb8 (TC IV) were previously shown to regulate the development of microbiota-specific pTreg cells within mesenteric lymph nodes (mLN) and large intestine (LI)^11,12^. In the small intestine, pTreg cells predominantly reflect food-antigen induced Treg cells^3,15^. To determine if TCs also regulate small intestinal pTreg generation, we examined individual siLN and small intestine lamina propria (siLP) in mice deficient in MHCII-restricted antigen presentation by RORγt^+^ cells (*MHCII^ΔRORγt^*) at 4 weeks of age. We observed a marked reduction in RORγt^+^Foxp3^+^ cells in all lymph nodes examined as well as Peyers Patches (PP) and siLP (**fig. S1B**). Histological analysis at 12 weeks of age demonstrated severe small intestinal inflammation with marked inflammatory cell infiltrate, epithelial hyperplasia and goblet cell loss with epithelial erosions (**fig. S1C** and **D**), confirming a critical role for RORγt^+^ APCs in preventing dysregulated immune responses within the small intestine. Since loss of small intestinal pTreg cells in *MHCII^ΔRORγt^* mice could also reflect loss of microbiota-specific pTreg cells, we tested the specific role of TCs in tolerance to food antigens by adoptive transfer of ovalbumin (OVA)–specific naïve OT-II CD4^+^ T cells into *MHCII^ΔRORγt^*mice at P14 followed by oral administration of OVA (**Fig. 1, C** and **D**). Administration of oral antigen in this time window coincides with first encounter with a host of dietary antigens upon weaning from breast-milk to solid food and reflects the physiological window for establishing tolerance to dietary proteins in mice. 7 days after transfer, ∼60% of OT-II cells in the siLN and siLP of *H2-Ab1^fl/fl^* wild-type mice expressed Foxp3 with equal proportions of RORγt^+^ and RORγt^−^ Foxp3^+^ cells (**Fig. 1, E** and **F**). In *MHCII^ΔRORγt^* mice, OT-II pTreg differentiation was almost completely abolished (**Fig. 1, F** and **fig. S1E**). Loss of pTreg differentiation was associated with enhanced OT-II Tbet^+^ T_H_1 cell differentiation (**Fig. 1, G** and **fig. S1F**). These results demonstrate a critical role for antigen presentation by RORγt^+^ APCs in food-specific pTreg differentiation.

**Figure 1.**
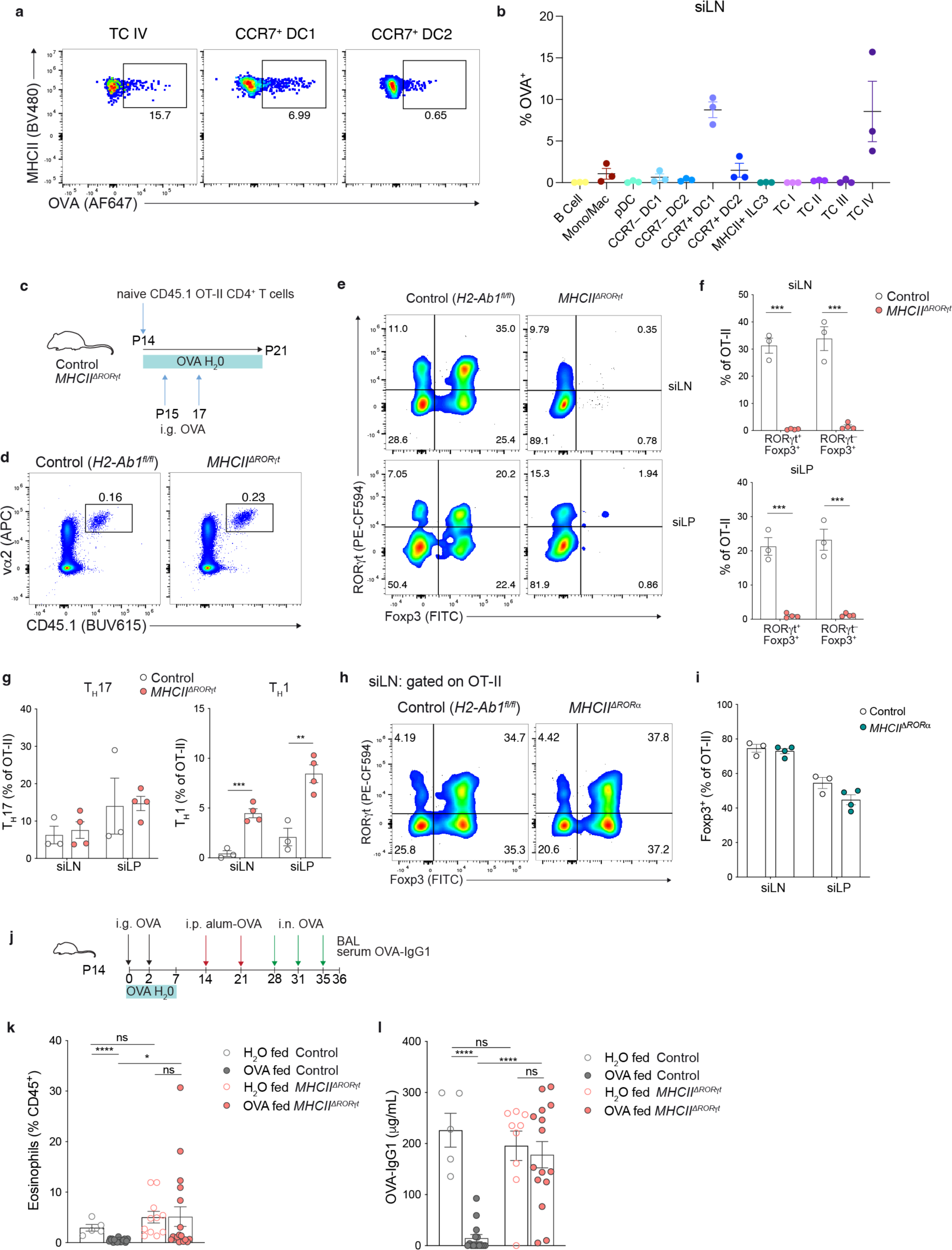
RORγt^+^ Thetis cells promote food-specific pTreg differentiation and oral tolerance. (**A**) Representative flow cytometry showing percentage of labeled TC IV, CCR7^+^ DC1 and CCR7^+^ DC2 in small intestinal mesenteric lymph nodes (siLN) of *Rorc^Venus-creERT2^*mice, 16 h after intra-gastric (i.g.) OVA-AF647 at postnatal day 14 (P14). (**B**) Summary graph of frequency of OVA-A647^+^ cells among indicated antigen-presenting cells (*n* = 3). (**C**-**G)** *MHCII^ΔRORγt^*(*n* = 4) and *H2-Ab1^fl/fl^* control (*n* = 3) littermate control mice were adoptively transferred with 25 x 10^3^ naïve CD45.1 CD4^+^ OT-II T cells prior to OVA administration and analyzed 7 d later for frequencies and numbers of RORγt^+^Foxp3^+^, RORγt^−^ Foxp3^+^ pTreg, T_H_1 (Foxp3^−^Tbet^+^) and T_H_17 (Foxp3^−^RORγt^+^) cells among OT-II cells in siLN and small intestine lamina propria (siLP). (**C**) Experimental set up. (**D**) Flow cytometry of CD45.1 TCRβ^+^vα2^+^ OT-II T cells in siLN. (**E**) Flow cytometry of RORγt and Foxp3-expressing OT-II cell subsets and (**F**) summary graphs for frequencies of OT-II pTreg cells. (**G**) Frequency of T_H_1 and T_H_17 cells among OT-II cells. (**H**) Flow cytometry of RORγt and Foxp3-expressing OT-II cell subsets and (**I**) summary graphs for frequencies of OT-II pTreg cells in siLN and siLP of *MHCII^ΔRORα^* (*n* = 4) and control (*H2-Ab1^fl/fl^*) (*n* = 3) mice. (**J**) Schematic of oral tolerance induction to alum-induced OVA immunity. (**K**) Eosinophil frequencies among CD45^+^ cells in broncho-alveolar lavage fluid from *H2-Ab1^fl/fl^*(control) or littermate *MHCII^ΔRORγt^* mice, with or without OVA feeding. (**L**) Serum OVA-IgG1 in mice described in **J**–**K**. Data in A, H, I representative of three independent experiments. Data in E-G representative of two independent experiments. Data in K and L pooled from 4 independent experiments. Error bars: means ± s.e.m. Each symbol represents an individual mouse. Statistics were calculated by two-tailed unpaired *t*-test; \**P* < 0.05; \*\**P* < 0.01; \*\*\**P* < 0.001; \*\*\*\**P* < 0.0001.

RORγt^+^ APCs encompass two distinct cell types – LTi-like type 3 innate lymphoid cells (ILC3) and TCs. Although we did not observe uptake of OVA in MHCII^+^ ILC3 cells (**Fig. 1B**), to definitively exclude a role for ILC3s in food-specific pTreg differentiation, we analyzed OT-II Treg conversion in *Rora^cre^H2-Ab1^fl/fl^*mice which specifically lack MHCII expression on ILC3s^11^. In contrast to *MHCII^ΔRORγt^* mice, OVA fed *MHCII^ΔRORα^* exhibited normal proportions and numbers of OT-II Treg cells (**Fig. 1, H** and **I**, and **fig. S1G**), excluding a role for ILC3s in food-specific pTreg generation.

To determine the impact of loss of food-specific pTreg cells on oral tolerance and Th2 allergic responses, *MHCII^ΔRORγt^* or control mice were immunized with OVA-Alum 7 and 14 days after the final exposure to oral OVA and challenged intranasally with OVA on three occasions, 3-4 days apart (**Fig. 1J**). Analysis of eosinophilia in the bronchoalveolar lavage fluid as well as OVA-specific IgG1, revealed suppression in OVA-fed wild-type but not *MHCII^ΔRORγt^* mice, relative to mice that received H_2_O gavages and regular drinking water (**Fig. 1, K** and **L**). Overall, these results indicate that food-specific Treg cells generated in early life comprise both RORγt^+^ and RORγt^−^ pTreg cells, and that MHCII antigen presentation by TCs is critical for food-specific pTreg generation and oral tolerance.

## Itgβ8^+^ TC IV are enriched at sites of pTreg generation in early life

We previously showed that CD11b^+^ TC IV express the TGF-β activating integrin αvβ8 and that Itgβ8 expression by TCs was required for colonic pTreg induction and tolerance to the microbiota^11^. To determine whether TC phenotypes were conserved across anatomically distinct lymph nodes, we performed scRNA-seq on TCs (CXCR6^−^RORγt^+^MHCII^+^) FACS-isolated from both gut and non-gut lymph nodes of *Rorc^Venus-creERT^*^2^ reporter mice at P14. Of note, although expression of Venus reflects transcription of the Rorc 3’-UTR, shared between both *Rorc* and *Rorgt* isoforms, *Rorgt^cre^* fate mapping confirmed that all TCs, including TC IV, express the RORγt isoform (**fig. S2A**). Differential gene expression analysis of TC cluster transcriptomes confirmed the presence of previously identified TC I-IV subsets (**Fig 2, A** and **B**). Analysis of TC composition across the different lymph nodes and tissues revealed a striking enrichment of TC IV within intestinal lymph nodes and gut associated lymphoid tissue including hepatic, small intestinal, colonic LNs and PPs (**Fig 2, C**). Amongst TCs, TC IV were distinguished by the highest levels of Itgb8 expression (**Fig 2, D**). In agreement with the scRNA-seq data, flow cytometry analysis of TCs from lymph nodes of P14 *Rorc^Venus-creERT2^Itgb8^tdtomato^* mice confirmed that the absolute number of Itgb8^+^ TC IV were increased in gut and hepatic lymph nodes relative to skin-draining lymph nodes (**Fig 2, E** and **F**). To determine if food-specific pTreg induction was dependent on Itgβ8 expression by TCs, we adoptively transferred naïve OT-II cells into P14 *Rorgt^cre^Itgb8^fl/fl^* mice followed by oral OVA administration. OT-II pTreg cell differentiation was abolished in these mice (**Fig 2, G** and **fig. S2B**). We did not observe altered OT-II pTreg generation in *Cd4^cre^Itgb8^fl/fl^*mice (**fig. S2C**), confirming that Itgβ8 expression by TCs, but not TCRβ^+^ cells, is required for food-specific pTreg induction.

**Figure 2.**
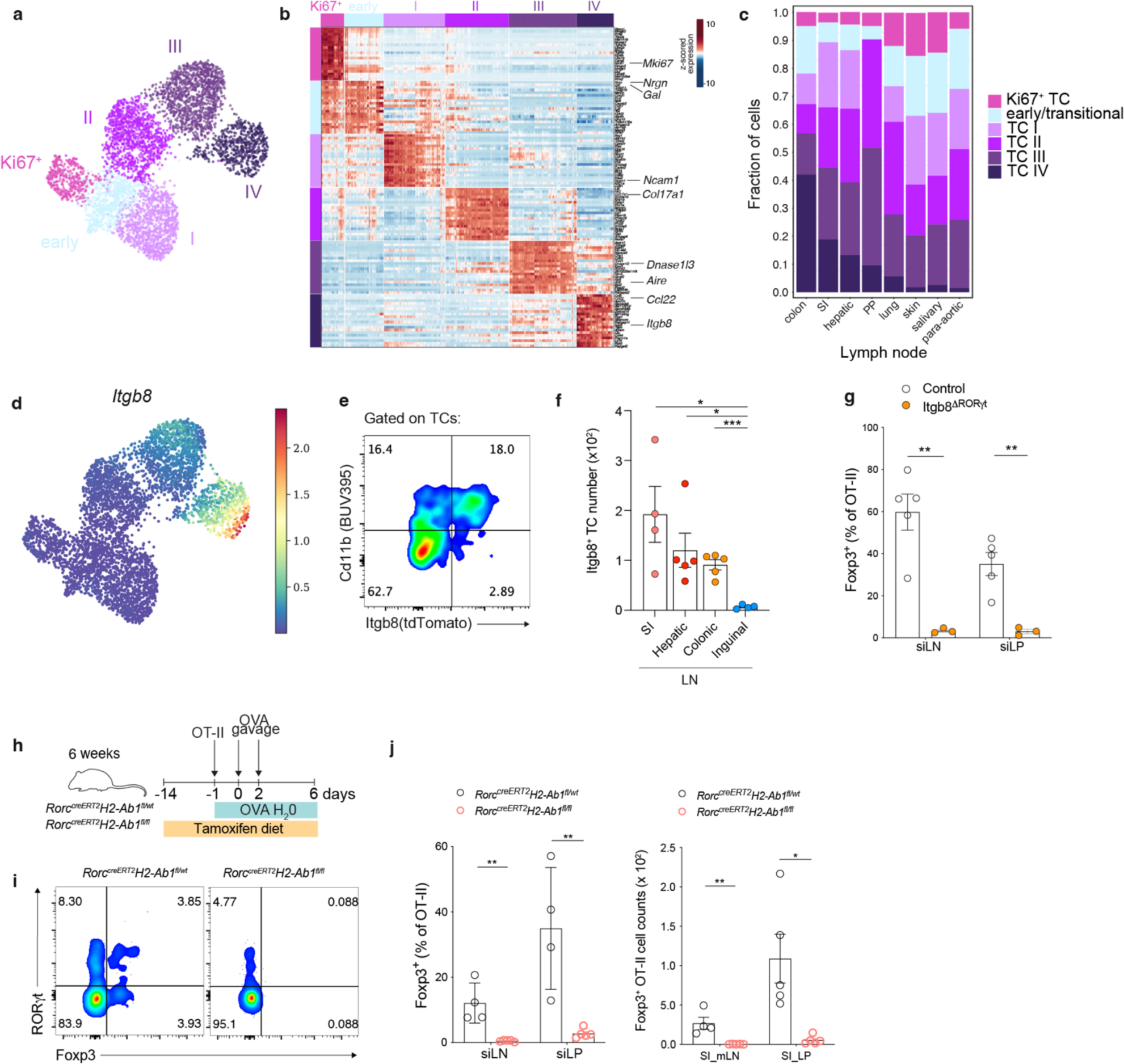
Enrichment of Itgb8^+^ TCs in early life gut lymph nodes provides a window of opportunity for enhanced food-specific pTreg generation. (**A**) Uniform manifold approximation and projection (UMAP) of 4,732 TCs from gut and non-gut lymph nodes of 2-week-old (P14) *Rorc^Venus-^*^c*reERT2*^ mice (*n* = 7), colored by cluster identity. (**B**) Heatmap showing expression of top differentially expressed genes between TC clusters identified in (A). (**C**) Fraction of each Thetis cell subset among all TCs in each lymph node sample. (**D**) UMAP overlaid by imputed expression of *Itgb8*. (**E**) Flow cytometry of Itgb8 expression in TCs isolated from mLN of P14 *Rorc^Venus-creERT2^Itgb8^tdTomato^*mice and (**F**) summary graph for total number of Itgβ8^+^ TCs in each indicated lymph node. (**G**) Frequencies of OT-II pTreg cells in siLN and siLP of *Itgb8^ΔRORγt^* (*n* = 3) and control (*Itgb8^fl/fl^*) (*n* = 5) mice. (**H**) Schematic for temporal deletion of MHCII, adoptive transfer of OT-II and OVA feeding in *Rorc^creERT2^H2-Ab1^fl/fl^* or littermate *Rorc^creERT2^H2-Ab1^fl/wt^* (control) mice. (**I**) Flow cytometry of RORγt and Foxp3-expressing OT-II cell subsets and (**J**) summary graphs for frequencies (left) and numbers (right) of OT-II pTreg cells in mice described in (H), *n* = 4 mice per group. Data in E and F representative of four independent experiments. Data in I and J representative of two independent experiments. Error bars: means ± s.e.m. Each symbol represents an individual mouse. Statistics were calculated by two-tailed unpaired *t*-test; \**P* < 0.05; \*\**P* < 0.01; \*\*\**P* < 0.001.

Given the previously reported developmental wave of TCs in mLNs^11^, we wondered whether the reduced number of TCs in adult mice was sufficient to promote food-specific pTreg cells following antigen introduction in later life, or whether a distinct antigen-presenting cell was responsible for food-specific Treg induction in adulthood. To address this, we continuously ablated MHCII expression in RORγt^+^ APCs in *Rorc^Venus-creERT2^H2-Ab1^fl/fl^* mice treated with tamoxifen from 6 weeks age, and adoptively transferred naïve OT-II T cells followed by oral OVA administration at 8 weeks (**Fig 2, H**). Analysis of OT-II cells 7 days later revealed an almost complete absence of OT-II pTreg cells in *Rorc^Venus-creERT2^H2-Ab1^fl/fl^*mice (**Fig 2, I** and **J**). Notably, however, the frequency of Foxp3^+^ cells among OT-II T cells in adult control mice was considerably less than that observed following OVA administration in mice at weaning age (12 vs 65%, siLN; **Fig 1E** and **2J**).

Together these results demonstrate that Itgb8^+^ TC IV promote pTreg induction throughout life; however, the increased abundance of TC IV in early life results in a window of opportunity for enhanced pTreg generation.

## Dendritic cells promote food-specific T_H_1 cells and maintain memory T cells

Our earlier experiments demonstrated uptake of OVA by both TC IV and CCR7^+^ DC1 (**Fig 1, A** and **B**). Previous studies suggested that tolerance to oral food antigens is mediated by conventional DCs, primarily DC1s^5^. To better understand the distinct roles of TCs and DCs in oral tolerance, we turned to *Clec9a^cre/cre^H2-Ab1^fl/fl^*(*MHCII^ΔDC^*) mice which specifically lack antigen presentation on DC1s, DC2s and pDCs^16^. Use of homozygous *Clec9a^cre^* mice allows efficient pan-DC targeting due to expression of Clec9a at the common DC progenitor (CDP) stage; however, neonatal DCs arise from an alternative progenitor with full transition to the classical Clec9a^+^ CDP differentiation pathway by 3 weeks of age^17^. We therefore adoptively transferred naive OT-II T cells into recipient mice at P21 followed by oral OVA administration (**Fig 3, A**). At this time-point, analysis of *Clec9a^cre/cre^R26^lsl-tdtomato^*mice confirmed effective labeling of both CCR7^−^ and CCR7^+^ DC1 and DC2 subsets (**Fig 3, B**). Similarly, we observed efficient ablation of MHCII expression on both XCR1^+^ DC1 and XCR1^−^ DC2s in siLN of *MHCII^ΔDC^* mice at this timepoint (**fig. S3A**). Analysis of OT-II cells 7 days after transfer revealed equivalent proportions of OT-II pTreg cells between *MHCII^ΔDC^* and littermate control *Clec9a^cre/cre^H2-Ab1^fl/wt^*mice (**Fig. 3, C**). In contrast, analysis of Foxp3^−^ effector T cells demonstrated impaired OT-II T_H_1 cell differentiation but equivalent proportions of T_H_17 cells (**Fig. 3, D**), suggesting that food antigen presentation by CCR7^+^ DC1s directs T_H_1 differentiation. However, analysis of OT-II cell numbers revealed a global reduction in the number of OT-II cells in siLN due to a loss of all Foxp3^+^ and Foxp3^−^ Teff subsets, including Teff cells lacking specific lineage markers (T_N_) (**Fig. 3, E**).

**Figure 3.**
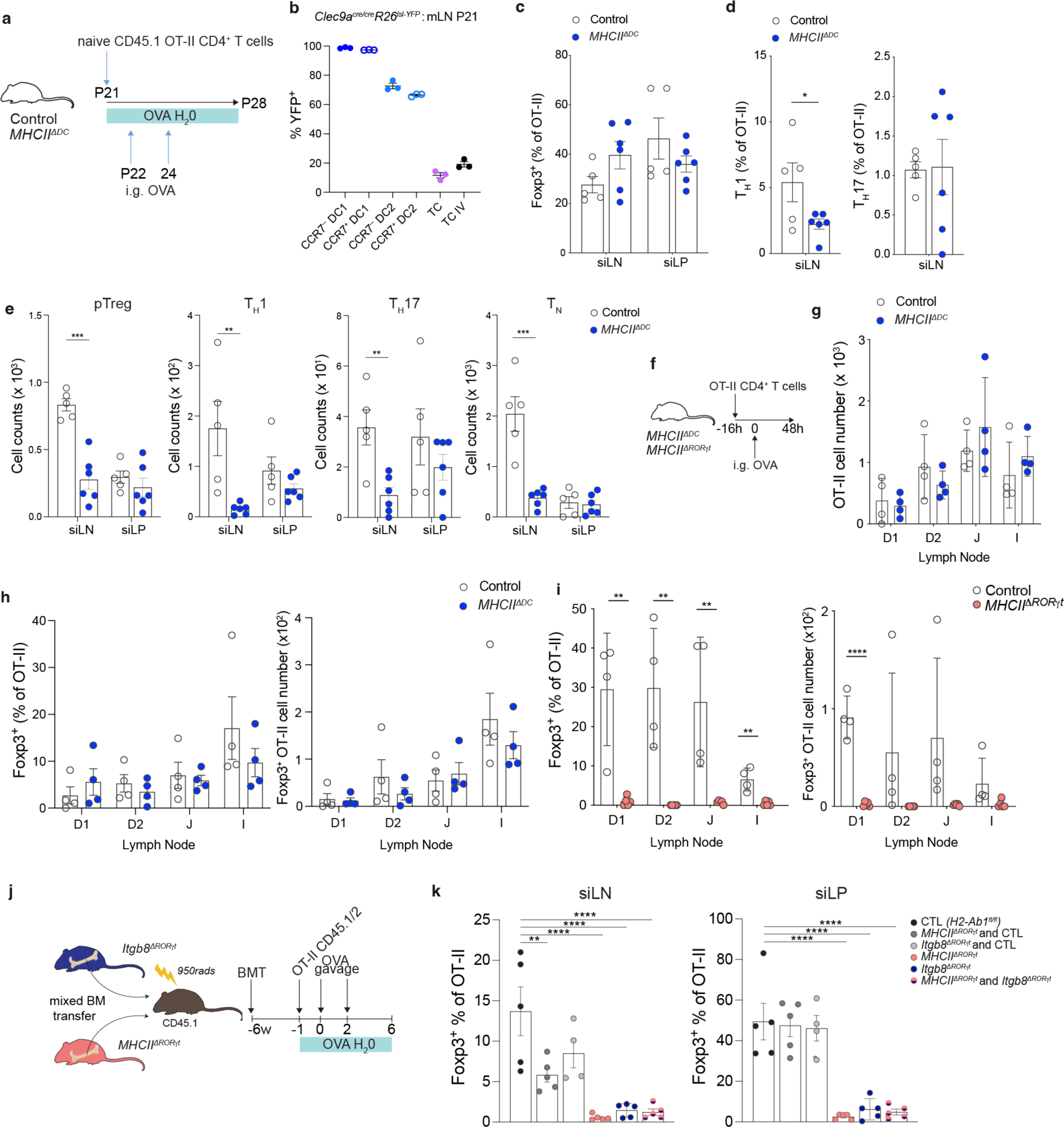
Antigen presentation by dendritic cells promotes food-specific T_H_1 differentiation and sustains memory T cells. (**A**) Schematic of OT-II adoptive transfers and OVA feeding. (**B**) Proportion of Clec9a fate-mapped (YFP^+^) cells in each indicated APC subset in mLN of P21 *Clec9a^cre/cre^R26^lsl-YFP^*mice. (**C-E**) Frequencies and numbers of OT-II pTreg cells, Foxp3^−^RORγt^−^Tbet^+^ T_H_1, Foxp3^−^RORγt^+^ T_H_17 and Foxp3^−^RORγt^−^Tbet^−^ T_N_ OT-II cells in siLN and siLP of *MHCII^ΔDC^* (*n* = 6) and control (*n* = 5) mice. (**F**) Schematic of OT-II adoptive transfers and i.g. OVA administration for analysis of pTreg induction 48hr post-OVA gavage. (**G**) Number of OT-II cells in individual siLN of *MHCII^ΔDC^* or control mice, *n* = 4 mice per group. (**H**) Frequency and number of OT-II pTreg cells in individual gut LN of *MHCII^ΔDC^* or control mice, *n* = 4 mice per group. (**I**) Frequency and number of OT-II pTreg cells in gut LN of *MHCII^ΔRORγt^* (*n* = 5) mice or control (*n* = 4) mice. (**J**). Mixed bone marrow chimera mice reconstituted with *MHCII^ΔRORγt^* and *Itgb8^ΔRORγt^* bone marrow cells were adoptively transferred with congenic OT-II T cells prior to oral OVA. (**K**) Frequency of Foxp3^+^ pTreg cells among OT-II cells in siLN and siLP of bone marrow chimeras. Data in B–E, I and J representative of three independent experiments. Data in G representative of two independent experiments. Gut lymph nodes: D1, portal; D2, distal duodenum; J, jejunum; I, ileum; C, colon. Error bars: means ± s.e.m. Each symbol represents an individual mouse. Statistics were calculated by two-tailed unpaired *t*-test; \**P* < 0.05; \*\**P* < 0.01; \*\*\**P* < 0.001, \*\*\*\**P* < 0.0001.

Given that our analysis of OT-II cells was performed after 7 days of continuous OVA ingestion via the drinking water, we wondered whether the reduction in numbers of antigen-experienced OT-II cells across all cell lineages was due to a role for DCs in maintaining effector-memory T cells through TCR signaling. To distinguish the role of TCs and DCs in induction vs maintenance of T cell effector-memory, we repeated the OT-II transfer experiments in both *MHCII^ΔDC^* and *MHCII^ΔROR^*^γ*t*^ mice and analyzed OT-II cells in individual gut draining lymph nodes, 48hrs after a single dose of i.g. OVA (**Fig. 3, F**). At this timepoint, OT-II cell numbers were normal in *MHCII^ΔDC^* mice, with equivalent proportions and numbers of Foxp3^+^ OT-II pTreg cells in all lymph nodes examined (**Fig. 3, G** and **H**). In contrast, OT-II Treg induction was completely abolished in *MHCII^ΔRORγt^* mice (**Fig. 3, I**). Together, these results indicate that antigen presentation by TCs but not DCs is required for initial priming and induction of pTreg cell differentiation. To further address potential redundancy by TCs and DCs in food-specific pTreg generation, we generated bone marrow chimeras with an equal mix of *Itgb8^ΔRORγt^* and *MHCII^ΔRORγt^* bone marrow cells—in which both DCs and TCs can present antigen, but the same TC cannot present antigen and activate TGF-β (**Fig. 3, J**). In these mice, OT-II pTreg induction was significantly impaired, with an equivalent deficit in OT-II pT_reg_ cells observed in *MHCII^ΔRORγt^*, *Itgb8^ΔRORγt^* and *MHCII^ΔRORγt^/Itgb8^ΔRORγt^* chimeras (**Fig. 3, K**), confirming i) that food-induced pT_reg_ differentiation is critically dependent on antigen presentation by Itgβ8-expressing TCs and ii) excluding the possibility of redundant and compensatory functions between TCs and DCs.

To further understand the role of DCs in maintenance of food-specific CD4^+^ T cells we analyzed cell proliferation in transferred OT-II cells, 72hrs after a single dose of i.g. OVA. At this timepoint, nearly all OT-II cells were CD44^hi^ and we did not observe differences in the proportion of CD44^hi^ cells in *MHCII^ΔDC^* mice (**fig. S3B** and **C)**. However, OT-II cell numbers were significantly reduced in individual gut LNs in *MHCII^ΔDC^* mice (**fig. S3D**). Comparison of cell proliferation revealed equivalent numbers of CD44^hi^ antigen experienced OT-II cells among undivided cells with progressive impairment in OT-II cell numbers with subsequent cell divisions (**fig. S3E** and **F)**, suggesting that antigen presentation by DCs is required to sustain early CD44^hi^ T cells.

Taken together, these results highlight distinct roles for TCs and DCs in regulation of food-specific T cell responses and suggest that antigen presentation and Itgβ8 expression by TC IV is required for induction of pTreg cells. In contrast, antigen presentation by DCs is required for food specific T_H_1 cell differentiation. Furthermore, our data highlight an additional role for antigen presentation by DCs in sustaining or amplifying antigen-experienced effector/memory T cells, irrespective of cell lineage.

## Discussion

T cell responses to food antigens are multifaceted, comprising both inflammatory effector T (Teff) cells and immunosuppressive pTreg cells. Prevention of dysregulated immune responses requires dominant suppression of Teff cells by pTreg cells. Thus, the balance between Treg and Teff cells is key to immune tolerance and intestinal homeostasis. Our study revealed a division of labor amongst intestinal APCs with recently identified TC IVs playing a critical role in pTreg generation, in contrast to induction of T_H_1 effector differentiation by DCs. Studies from both mouse and humans have established a window of opportunity for enhanced oral tolerance in early life. Our findings show that the balance of Treg/Teff cells is skewed heavily in favor of Treg cell differentiation following antigen-encounter in the peri-weaning window, a period associated with increased abundance of tolerogenic TCs within gut lymph nodes.

Unexpectedly, our studies revealed a role for DCs in maintaining activated/memory Teff and Treg cells following lineage fate commitment. A wealth of studies examining CD8 T cell differentiation have demonstrated the development of TCF1^hi^ ‘stem-like’ memory T cells that sustain CD8 effectors in settings of chronic antigen availability^18^. Analogous populations of CD4 stem cell memory T cells have recently been identified in lymph nodes during transplantation, tumorigenesis and autoimmunity^19–22^. Since continuous dietary exposure provides chronic antigen stimulation, it seems likely that food-specific T cells would be sustained by such progenitors. Our findings suggest that antigen presentation by dendritic cells may play a key role in sustaining lymph node stem-like memory T cells or their progeny and will be an important area of future investigation.

The discovery that TC IVs regulate tolerance to food as well as gut microbiota, suggests a broad role for TCs in intestinal tolerance to harmless foreign antigens. The recent characterization of TCs in humans^11,23^, alongside our discovery that TCs instruct intestinal pTreg differentiation both in early life and adulthood, suggests potential therapeutic value in targeting antigens to TCs for promotion of intestinal immune tolerance in settings of food allergy and intestinal inflammatory diseases.

## Acknowledgements

We thank the Single-cell Analysis and Innovation Lab (SAIL) at MSK for sample processing, Helena Paidassi for provision of *Itgb8^tdTomato^* mice and Michael Glickman for OT-II mice. We also thank Maria Canesso and Daniel Mucida for discussions and advice. We acknowledge the use of the Integrated Genomics Operation Core, funded by the NCI Cancer Center Support Grant (CCSG, P30 CA08748), Cycle for Survival, and the Marie-Josée and Henry R. Kravis Center for Molecular Oncology.

## Author Contributions

Y.F.P. and V.C. designed experiments, analyzed data and edited the manuscript; T.P. designed and performed computational analyses; B.A., Z.Z., Y.L., G.S., Y.P.I. and L.F. performed experiments; C.L. supervised computational analyses. C.C.B. conceived, designed and supervised the research. All authors read and approved the manuscript.

## Data and materials availability

The sequencing data are available through the Gene Expression Omnibus under accession GSE (pending).

## Materials and Methods

### Mice

*Rorc^Venus-T2A-creERT2^*, *Clec9a^cre^, Rora^cre^, Itgb8^fl/fl^, Cd4^cre^, OT-II* and *Itgb8^tdTomato^*mice have been previously described^11,16,24–28^. *Rorgt^cre^, H2-Ab1^fl/fl^, R26^lsl-tdTomato^, R26^lsl-YFP^*, C57Bl/6 (CD45.2), and CD45.1 mice were purchased from Jackson Laboratories. Generation and treatments of mice were performed under protocol 21-05-007, approved by the Sloan Kettering Institute (SKI) Institutional Animal Care and Use Committee. All mouse strains were maintained in the SKI animal facility in specific pathogen free (SPF) conditions in accordance with institutional guidelines and ethical regulations. Both male and female mice were included in the study and we did not observe sex-dependent effects. All mice analyzed were age and litter matched unless otherwise specified. All animals used in this study had no previous history of experimentation and were naïve at the time of analysis.

#### Tamoxifen diet

*Rorc^Venus-creERT2^H2-Ab1^fl/fl^*and littermate *Rorc^Venus-creERT2^H2-Ab1^fl/fwt^*mice were placed on a tamoxifen citrate–containing diet (TD.130860; Envigo) at 6 weeks of age for 2 weeks prior to adoptive T cell transfer and for the duration of the experiment.

### Tissue processing

Mice were euthanized by CO_2_ inhalation. Organs were harvested and processed as follows. Lymphoid organs were digested in collagenase in RPMI1640 supplemented with 5% fetal calf serum, 1% L-glutamine, 1% penicillin–streptomycin, 10 mM HEPES, 1 mg/ml collagenase A (Sigma, 11088793001) and 1U/mL DNase I (Sigma, 10104159001) for 45 min at 37°C, 250 rpm. Small intestine lymph nodes compromised the main mesenteric lymph node chain with the colon lymph node removed. For analysis of individual gut lymph nodes, lymph nodes draining intestinal segments were identified anatomically^29^. Large and small intestines were removed and PPs were removed from small intestines. Intestines were flushed with PBS and incubated in PBS supplemented with 5% fetal calf serum, 1% L-glutamine, 1% penicillin–streptomycin, 10 mM HEPES, 1 mM dithiothreitol, and 1 mM EDTA for 15 min to remove the epithelial layer. Samples were washed and incubated in digest solution for 30 min. 1/4 inch ceramic beads (MP Biomedicals, 116540034) were added to intestine samples (3 per sample) to aid in tissue dissociation. Digested samples were filtered through 100-μm strainers and centrifuged to remove collagenase solution.

### Flow cytometry

For flow cytometric analysis, dead cells were excluded by staining with LIVE/DEAD Fixable Zombie NIR in PBS for 10 minutes at 4°C, prior to cell-surface staining, along with anti-CD16/32 to block binding to Fc receptors. Extracellular antigens were stained for 30 minutes at RT in Brilliant Violet staining buffer (BD Biosciences). For intracellular protein analysis, cells were fixed and permeabilized with Cytofix (BD Biosciences) or Ebioscience Foxp3 kit, per manufacturer instructions. Intracellular antigens were stained for 30min at 4°C in the respective 1x Perm/Wash buffer. Cells were washed with staining buffer prior to acquisition on a Cytek Aurora™. The antibodies used for flow cytometry and FACS are listed in **Supplementary Table 1**. Unless otherwise stated, we used the following gatings: TCs: Lin (Siglec-F, TCRβ, TCRγδ, CD19, B220, NK1.1)^−^CD88^−^Ly6C^−^RORγt^+^ (intracellular staining or expression of Venus in *Rorc^Venus-creERT2^*mice)CXCR6^−^MHCII^+^; MHCII^+^ ILC3s: Lin^−^CD88^−^Ly6C^−^RORγt^+^CXCR6^+^MHCII^+^, and DCs: Lin^−^ CD88^−^Ly6C^−^RORγt^−^CD11c^+^MHCII^+^.

#### OVA antigen uptake assay

To evaluate OVA uptake, OVA (grade III, Sigma, A5378) was conjugated to Alexa Fluor-647 dye using Alexa Fluor™ 647 NHS Ester kit (Invitrogen) as per manufacturer’s instructions. P14 *Rorc^Venus-creERT2^*mice were gavaged with 200μl of 20mg/ml OVA-AF647 and small intestine mesenteric lymph nodes were analyzed 16 h later.

### Histological analysis of intestinal inflammation

Mice were euthanized by CO_2_ inhalation and small intestines were harvested and immediately placed into 10% formalin. Histopathological assessment for inflammation scoring in the intestine was performed on H&E stained sections based on established scoring systems for intestinal inflammation in mouse models^30^. Assessment includes severity and extent of inflammatory cell infiltrates, epithelial changes and mucosal architecture changes. Briefly, severity and extent of inflammatory cell infiltrate in the mucosa and if extending to submucosa and muscularis were evaluated histologically. Other evaluations include proliferation of epithelial cells lining the mucosa villous atrophy, crypts, loss of goblet cells, crypt abscesses, erosions and ulceration.

#### Adoptive T cell transfer

CD4^+^ T cells were pre-enriched from spleens of CD45.1 or CD45.1/2 OT-II mice by negative selection using the CD4^+^ T-cell isolation kit (Miltenyi Biotec). Live (SytoxBlue^−^)CD25^−^ CD44^lo^CD62L^+^TCRβ^+^vα2^+^ naïve OT-II T cells were FACS-isolated. For adoptive transfer of OT-II T cells and analysis 7 days post-transfer, 2.5 x10^3^ (P14 recipients), 5 x10^3^ (P21 recipients) or 7.5 x 10^5^ (adult recipients) cells were transferred by retro-orbital injection. For short-term analysis of OT-II T cells (48 or 72hrs post-OVA gavage), mice received 2.5 x 10^5^ cells by retro-orbital injection.

#### Analysis of OT-II T cell proliferation

FACS-isolated naïve CD45.1 OT-II T cells were labelled using CellTrace Violet Cell Proliferation Kit (Invitrogen C34557) as per manufacturer’s instructions. 2.5 x 10^5^ labelled naïve OT-II T cells were transferred into P21 recipient mice by retro-orbital injection. Mice were intra-gastrically gavaged with 25mg OVA (grade III, Sigma, A5378) in 100μL, 16 h post adoptive cell transfer and analyzed 72 hours post-gavage.

#### Oral antigen administration

OVA (grade III, Sigma, A5378) was administered intragastrically using plastic gavage needles. P14-21 mice were gavaged with 25mg in 100μl of H_2_O, 1 and 3 days after adoptive transfer of OT-II T cells and simultaneously fed *ad libitum* with OVA dissolved in drinking water (0.75% w/v). Adult mice were gavaged with 100mg of OVA in 200μl of H_2_O and fed *ad libitum* with OVA dissolved in drinking water (1.5% w/v).

#### Alum immunization and airway challenge

14 and 21 days after first oral administration of OVA, 40mg of endotoxin free OVA antigen (Invivogen) adsorbed to 40μl of Alum hydrogel (Invivogen) was injected intraperitoneally (i.p.) in a final volume of 200μl PBS. 14, 17 and 21 days after the first i.p. immunization, mice were anaesthetized and 10μg of endotoxin-free OVA (Invivogen) was administered intranasally in 35μl of PBS.

#### Bronchoalveolar lavage fluid (BALF) analysis

Mice were euthanized, the trachea was cannulated, and lungs were washed with 1.0ml of sterile PBS. Total BALF cells were stained for flow cytometry analysis.

#### Anti-OVA IgG1 ELISA

Serum OVA-specific IgG1 was detected using Anti-Ovalbumin IgG1 (mouse) ELISA Kit (Cayman Chemical 500830) following the manufacturer’s protocol.

### scRNA-sequencing

For single cell RNA seq of TCs, lymph nodes from P14 *Rorc^Venus-creERT2^* mice were pooled from 7 biological replicates and processed as described earlier. Cells were depleted of Lineage (TCRb, TCRγ8, CD19, B220, NK1.1)^+^ cells via staining with biotinylated antibodies followed by magnetic bead negative selection. Cells were incubated with anti-CD16/32 in sorting buffer (2% FBS in PBS) for 10 minutes at 4°C to block binding to Fc receptors. Extracellular antigens were stained for 30 minutes at 4°C in sorting buffer (2% FBS, 2mM EDTA, in PBS). Cells from distinct lymph nodes were “stained” with TotalSeq^TM^ oligo-conjugated antibodies consisting of a pool of antibodies directed against CD45 (30-F11) and H2 MHC Class I alloantigens (clone M1/42; Biolegend). Cells were washed and resuspended in sorting buffer with SYTOX blue (Invitrogen) for exclusion of dead cells. Live, CD45^+^Lin(Siglec-F, TCRý, TCRy8, CD19)^−^ RORyt(Venus)^+^CXCR6^−^MHCII^+^ cells were then sort purified. Cells were sorted into cRPMI, before being pelleted and resuspended in RPMI-2% FBS to a final concentration of 700–1,300 cells per μl. Single Cell Gene Expression was performed on Chromium instrument (10X Genomics) following the user guide manual for 3′ v3.1. In brief, viability of cells was confirmed above 80% with 0.2% (w/v) Trypan Blue staining (Countess II). Cells were captured in droplets. Following reverse transcription and cell barcoding in droplets, emulsions were broken and cDNA purified using Dynabeads MyOne SILANE followed by PCR amplification per manual instruction. ∼10,000 cells were targeted in total. Cells from different lymph nodes were multiplexed together on one lane of 10X Chromium using Hash Tag Oligonucleotides - HTO following previously published protocol^31^. Final libraries were sequenced on Illumina NovaSeq S4 platform (R1 – 28 cycles, i7 – 8 cycles, R2 – 90 cycles).

### Mouse single-cell RNA-seq computational analysis

#### Pre-processing of the 10X scRNA-seq for Thetis cells

Single-cell RNA-seq FASTQ files were aligned to mm10 (Cell Ranger mouse reference genome mm10-2020-A) and counted by Cell Ranger v7.1.0 with default parameters. The barcodes were filtered based on the number of RNA-seq transcripts (>1,500), the number of detected genes (>500), and the fraction of mitochondrial transcripts (<5%). Finally, any genes detected in <2 cells in the scRNA-seq data were discarded, leaving 19,666 genes. After clustering the scRNA-seq data (described in ‘Dimensionality reduction, cell clustering, and visualization’), and based on the expression of marker genes, we identified 3 contaminant clusters (CXCR6^+^ ILC3: cluster 7, DC: cluster 8, low quality: cluster 9) which were excluded from downstream analyses. In total, 4,732 cells remained, with a median scRNA-seq library-size of 10,504.

#### Dimensionality reduction, cell clustering, and visualization

The filtered count matrix was library-size normalized, log-transformed (‘log-normalized’ expression values) and then centered and scaled (‘scaled’ expression values) using Seurat v4.4.0. Principal component analysis (PCA) was performed on the scaled data (npcs=50). A nearest-neighbor graph was constructed using the first 30 principal components (PCs) with 30 nearest neighbors. Clustering was performed using Louvain algorithm with resolution 0.6 on the shared nearest-neighbor graph. Cell clustering was visualized using UMAP^32^, computed from the same nearest neighbor graph used for clustering.

#### Differential gene expression tests

Differentially expressed genes (DEGs) between groups of cells were identified with MAST^33^, performed using Seurat functions. MAST was run on the log-normalized expression values. In all tests, genes were only considered if they were detected in at least 1% of the cells in at least one of the two groups compared (min.pct=0.01, logfc.threshold=0). In one-vs-rest DE tests comparing multiple groups, each group was compared to all the cells from other groups. Specific DE comparisons are described in the results. DEGs were reported according to their log-fold change (>1.5) and adjusted *p*-value (<0.01). Ribosomal and mitochondrial genes were removed from the final list of genes reported/visualized. Where stated, the top DEG markers were subsequently selected for each group, based on fold change.

#### Data imputation for scRNA-seq data

MAGIC imputation^34^ was applied to the log-normalized expression values to further de-noise and recover missing values. Imputed gene expression values were only used for data visualization on UMAP overlays and heatmaps, where stated.

### Bone marrow chimera mice

Bone marrow (BM) cells were isolated from indicated donor mice, depleted of CD90.2^+^ and TER-119^+^ cells using magnetic bead-based depletion. BM cells were resuspended in PBS and 2-3 x 10^6^ cells were injected into 6-week-old CD45.1 mice, irradiated with 950rads/mouse one day earlier.

### Statistical analysis

Analysis of all data was done with unpaired two-tailed t test with a 95% confidence interval, as specified in the text or legends. *P* < 0.05 was considered significant: * *p* < 0.05; ** *p* < 0.01; \*\*\**p* < 0.001; \*\*\*\**p* < 0.0001. Details as to number of replicates, sample size, significance tests, and value and meaning of *n* for each experiment are included in the Methods or Figure legends. Statistical tests were performed with Prism (GraphPad Software). scRNA-sequencing experiments were carried out once. Mice were non-randomly allocated to experimental groups to ensure equal distribution of genotypes between treatments. Researchers were not blinded as to genotype or treatment during the experiments. No measures were taken to estimate sample size of to determine whether the data met the assumptions of the statistical approaches used. Significance (α) was defined as < 0.05 throughout, after correcting for multiple comparisons.

### Data and material availability

The mouse sequencing data are available through the Gene Expression Omnibus under accession number (pending).

---

**Fig. S1.**
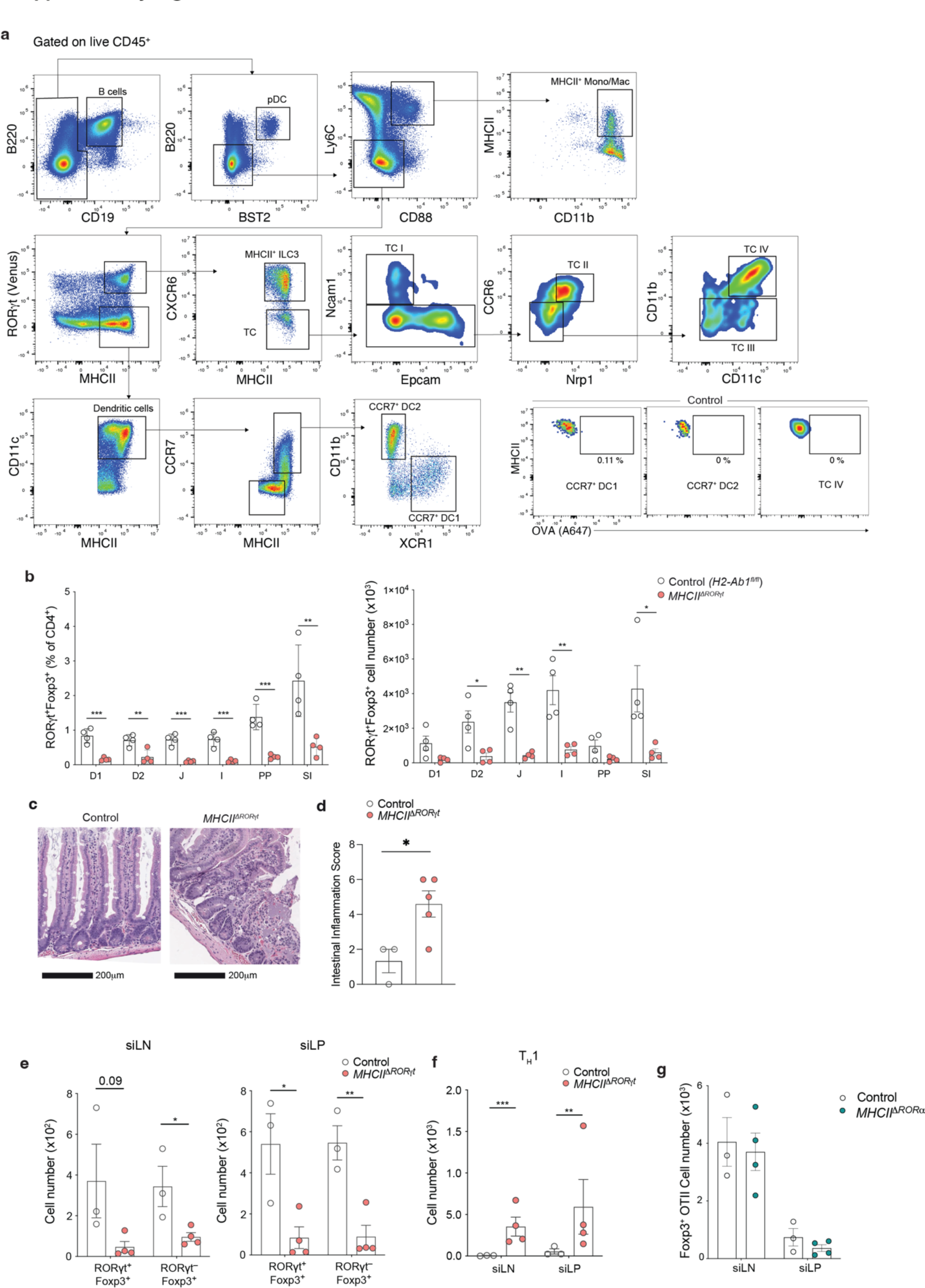
RORγt^+^ Thetis cells promote food-specific pTreg differentiation and oral tolerance. (**A**) Representative flow cytometry showing gating of MHCII^+^ antigen-presenting cell types in small intestinal lymph nodes (siLN) of *Rorc^Venus-creERT2^*mice 16 h after intra-gastric (i.g.) OVA-AF647 at postnatal day 14 (P14). (**B**) Frequency and number of RORγt^+^Foxp3^+^ pTreg cells in individual gut lymph nodes and small intestine lamina propria (SI) of *MHCII^ΔRORγt^* and *H2-Ab1^fl/fl^* littermate mice (*n* = 4 mice per group) at 4 weeks of age. (**C**) Representative haematoxylin and eosin (H&E) stained sections of the small intestine from *MHCII^ΔRORγt^* and control mice at 12 weeks of age. Scale bars, 200 μm. (**D**) Histological small intestinal inflammation score in 12-week-old *MHCII^ΔRORγt^* (*n* = 5) and control (*n* = 3) mice. (**E**) Number of RORγt^+^Foxp3^+^ and RORγt^−^Foxp3^+^ OT-II pT_reg_ cells and (**F**) Foxp3^−^RORγt^−^Tbet^+^ T_H_1 OT-II cells in siLN and small intestine lamina propria (siLP) of *MHCII^ΔRORγt^* (*n* = 4) and control (*H2-Ab1^fl/fl^*) (*n* = 3) mice. (**G**) Number of Foxp3^+^ OT-II pTreg cells in siLN and siLP of *MHCII^ΔRORα^* (*n* = 4) and control (*H2-Ab1^fl/fl^*) (*n* = 3) mice. Data in A, B and G representative of three independent experiments. Data in C from one experiment. Data in E-F representative of two independent experiments. Gut lymph nodes: D1, portal; D2, distal duodenum; J, jejunum; I, ileum; C, colon. Error bars: means ± s.e.m. Each symbol represents an individual mouse. Statistics were calculated by two-tailed unpaired *t*-test; \**P* < 0.05; \*\**P* < 0.01; \*\*\**P* < 0.001; \*\*\*\**P* < 0.0001.

**Fig. S2.**
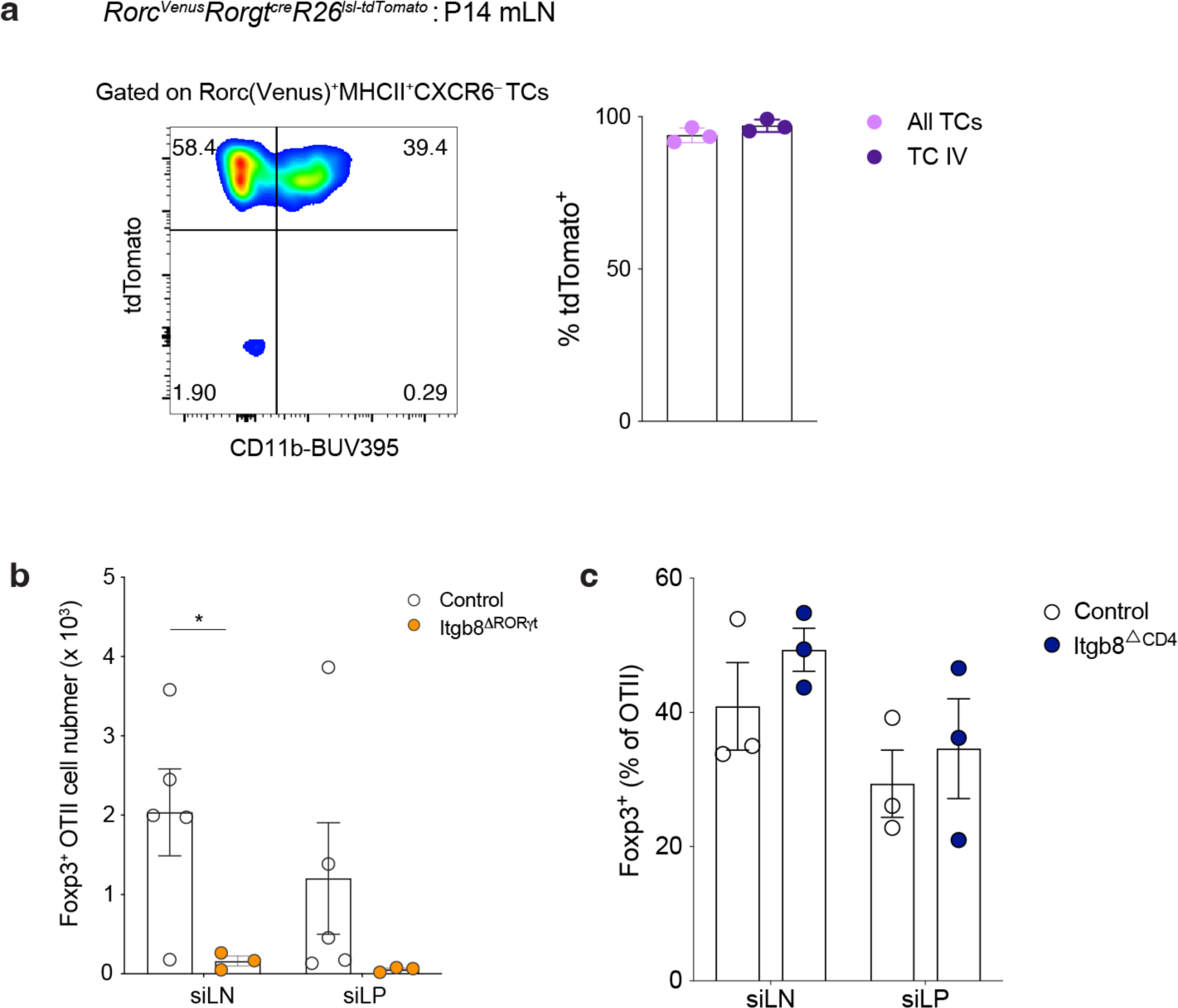
Itgb8-dependent Induction of food-specific pTreg cells by TC IV. (**A**) Flow cytometry of Lin^−^RORγt(Venus)^+^CXCR6^−^MHCII^+^ TCs from mLN of P15 *Rorgt^cre^R26^lsl-tdtomato^Rorc^Venus-CreERT2^*mice and summary graph of frequency of tdTomato^+^ cells amongst all TCs and CD11b^+^ TC IV (*n* = 3 mice). (**B**) Number of Foxp3^+^ OT-II pTreg cells in siLN and siLP of *Itgb8^ΔRORγt^* (*n* = 5) and control (*Itgb8^fl/fl^*) (*n* = 3) mice. (**C**) Number of Foxp3^+^ OT-II pTreg cells in siLN and siLP of *Itgb8^ΔCD4^*and control (*Itgb8^fl/fl^*) mice (*n* = 3 mice per group). Data in A and C representative of two independent experiments. Data in B representative of three independent experiments. Error bars: means ± s.e.m. Each symbol represents an individual mouse. Statistics were calculated by two-tailed unpaired *t*-test; \**P* < 0.05.

**Fig. S3.**
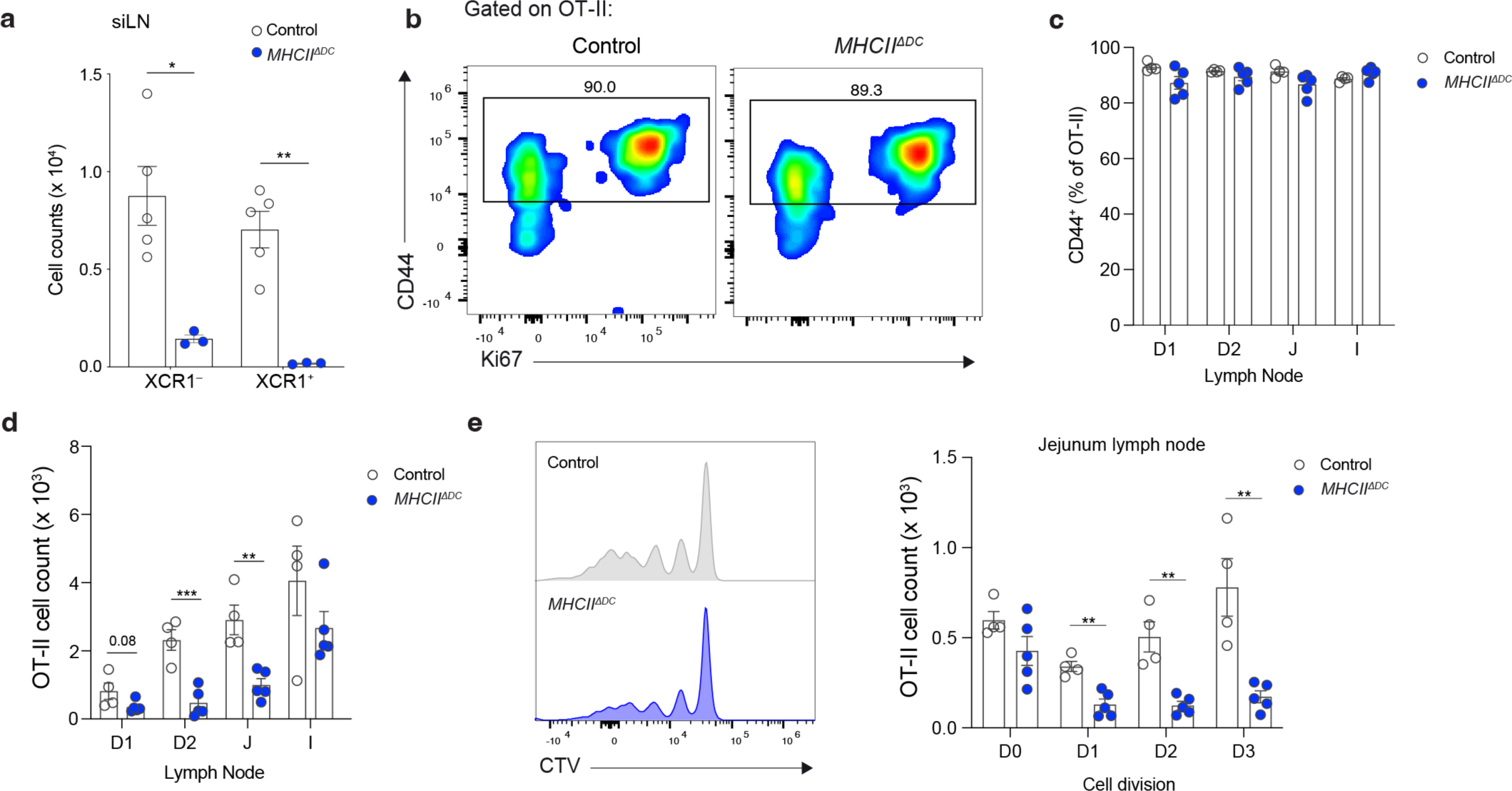
Antigen presentation by dendritic cells sustains memory T cells. **(A)** Number of dendritic cells in siLN of *Clec9a^cre/cre^H2-Ab1^fl/fl^* (*MHCII^ΔDC^*) and *Clec9a^cre/cre^H2-Ab1^fl/wt^* (control) mice at P21. Schematic of OT-II adoptive transfers and OVA feeding. (**B**) Representative flow cytometry of OT-II T cells in J-lymph node, 72 h after i.g. OVA. (C) Percentage of CD44^hi^ cells among OT-II T cells in gut lymph nodes, 72 h after one dose of i.g. OVA. (**D**) Total number of OT-II T cells in gut lymph nodes, 72 h after one dose of i.g. OVA. (**E**) Representative flow cytometry of OT-II cell proliferation and summary graph of number of J-LN OT-II T cells within each cell division; D0 = undivided. Data in A-E representative of two-three independent experiments. Error bars: means ± s.e.m. Each symbol represents an individual mouse. Statistics were calculated by two-tailed unpaired *t*-test; \**P* < 0.05; \*\**P* < 0.01; \*\*\**P* < 0.001.

